# Modulation of the mTOR pathway plays a central role in dendritic cell functions after Echinococcus granulosus antigen recognition

**DOI:** 10.1101/2020.07.27.224055

**Authors:** Christian Rodriguez Rodrigues, María Celeste Nicolao, Maia Chop, Natalia Plá, Mora Massaro, Julia Loos, Andrea C. Cumino

## Abstract

Immune evasion is a hallmark of persistent echinococcal infection, comprising modulation of innate immune cells and antigen-specific T cell responses. However, recognition of *Echinococcus granulosus* by dendritic cells (DCs) is a key determinant of the host’s response to this parasite. Given that mTOR signaling pathway has been described as a regulator linking metabolism and immune function in DCs, we reported for the first time in these cells, global translation levels, antigen uptake, phenotype, cytokine transcriptional levels, and splenocyte priming activity upon recognition of the hydatid fluid (HF) and the highly glycosylated laminar layer (LL). We found that LL induced a slight up-regulation of CD86 and MHC II in DCs and also stimulated the production of IL-6 and TNF-α. By contrast, HF did not increase the expression of any co-stimulatory molecules, but also down-modulated CD40 and stimulated the expression of the anti-inflammatory cytokine IL-10. Both parasitic antigens promoted protein synthesis through mTOR activation. The use of rapamycin decreased the expression of the cytokines tested, empowered the down-modulation of CD40 and also reduced splenocyte proliferation. Finally, we showed that *E. granulosus* antigens increase the amounts of LC3-positive structures in DCs which play critical roles in the presentation of these antigens to T cells.

## 2. Introduction

The cestode *Echinococcus granulosus* (*Eg*) is the etiological agent for cystic echinococcosis. This parasite infects humans as an intermediate host in a zoonotic transmission cycle with worldwide distribution ^1,2^. Cystic echinococcosis is considered as a re-emerging and neglected disease that causes serious chronic liver pathology, with a high fatality rate and poor prognosis without careful clinical management. In the viscera of the intermediate host, the parasites develop fluid-filled cysts, called hydatid cysts or metacestodes, that are isolated from the host by three layers. From inside to outside, an internal germinal layer composed of totipotent cells, with a high metabolic rate and in continuous cell division, that generates brood capsules and protoscoleces constituting the “hydatid sand” ^3^. Furthermore, the parasite synthesizes the laminar layer (LL), an acellular structure rich in mucins and polysaccharides that surrounds the metacestode, creating a mechanical and immunological protective barrier, crucial in the *Echinococcus*–host interface ^4,5^. Externally, and as a product of the host’s response, an adventitial layer, composed of newly formed eosinophils, fibroblasts, M2 macrophages, and capillaries, surrounds the LL. Under pressure and inside of the cyst it is found the hydatid fluid (HF) constitute by different macromolecules including a wide range of parasite and host proteins ^6^. During the infection course, the cyst may rupture or slowly leak out, permeate into a coelomic cavity, open on an epithelial surface or empty into the lumen of a hollow organ, leading to different host immune responses, ranging from a severe anaphylactic shock to a spontaneous resolution of the infection ^7^. In this context, *Eg-*antigens of metacestodes exposed and released in the tissues, lymphatics and circulation could be detected by dendritic cells (DCs).

Dendritic cells as antigen-presenting cells play a key role in the immune homeostasis, but under stress conditions rapidly respond to local or systemic perturbations by pathogenic or sterile insults ^8^. DCs capture antigens in the periphery or in the blood and migrate to the lymphoid organs, where they activate the specific immune response by inducing cytokine release and presenting processed peptides in a MHC molecule to specific T cells ^9^. This activation process, that includes cellular shape changes, migration, expression of co-stimulatory molecules in the membrane, production of cytokines, chemokines, lipid mediators, and antigen presentation must be metabolically supported to be efficient.

The mammalian target of rapamycin (mTOR) signaling pathway serves as a central regulator of cell metabolism, growth, proliferation and survival. mTOR is a serine/threonine protein kinase that forms the catalytic subunit of two distinct protein complexes, known as mTOR Complex 1 (mTORC1) and 2 (mTORC2) ^10^. These protein complexes sense growth factors, nutrients, and energy cell status ^11^. While mTORC1 plays a key anabolic role in promoting cell growth and proliferation, mTORC2 influences cell morphology through the regulation of cytoskeletal organization ^12^. Activation of the phosphatidylinositol 3-kinase (PI3K)– protein kinase B (AKT)– mTORC1 pathway is required for the *in vivo* development of normal numbers of mouse DCs in the presence of Flt3L ^13^. In recent years, the reports that link the immunological pathways of microbial sensing through different pattern recognition receptors to mTOR-dependent metabolic pathways have increased ^14–17^. In fact, the bacterial lipopolysaccharide improved global protein synthesis via TLR4-activation in human monocyte-derived DCs ^18^. Additionally, treatment of human and mouse DCs or monocytes with rapamycin enhanced their production of IL-12p40 and IL-12p70 and strongly reduced the expression of the anti-inflammatory cytokine IL-10 after stimulation with TLR ligands ^13,19–25^. Moreover, inhibition of mTOR abolished the ability to induce IFN-α and pro-inflammatory cytokine production by plasmacytoid dendritic cells ^26,27^, and promoted in DCs the expression of the co-stimulatory molecule CD86, whereas PD-L1, a negative regulator for T cell activation, is decreased ^19,22,25,28^. In addition, C-type lectin receptors recognized complex glycan structures in pathogens and modulated the immune cell functions by mechanisms dependent on the activation of mTOR ^29^. Rapamycin was also described to augments autophagy, which is important for the presentation of endogenous and exogenous proteins on MHC class I and class II molecules, thereby promoting activation of CD8^+^ and CD4^+^ T-cells respectively ^30^. On the other hand, mTOR modulation is also a target by pathogens to improve their survival. In *Leishmania* the protease GP63 cleaves mTOR on macrophages, inhibiting the mTORC1 with the concomitant activation of the translational repressor 4E-BP1 and consequently promoting parasite proliferation ^31^. It has been reported that upon infection with *Toxoplasma gondii*, PI3K/Akt/mTOR pathway is activated in macrophages, and this activation is required to avoid host cell defense ^32^. In helminths, *Brugia malayi* inhibits the phosphorylation of mTOR and its downstream proteins promoting autophagy ^33^ The hookworm *Nippostrongylus brasiliensis* modulates mTOR pathway in macrophages to induce differentiation and functional abilities of M2 profile and *Schistosoma mansoni*-soluble egg antigens via mTOR-dependent and -independent pathway condition human DCs to skew immune responses towards Th2 response ^34,35^.

Recently, the modulation of these metabolic pathways by the parasite *E. granulosus* has been reported. *Eg* excretory/secretory products activate the PI3K/AKT/mTOR pathway and the recruitment of alternatively activated macrophages ^36^. Also, *Eg* LL inhibits macrophage and CD11c^+^ antigen presenting-cells (APCs) proliferation in response to IL-4 and M-CSF *in vivo* and *in vitro* ^37^. Furthermore, the upregulation of the co-stimulatory molecule CD40 was inhibited in DCs by interfering with Akt and GSK3 activation ^38^. In the present manuscript, we address how the recognition of purified laminar layer (pLL) or HF from *E. granulosus* by DCs, induces phenotypical and functional changes in a mTORC1-dependent manner, with a subsequent *in vitro* splenocyte proliferation required to trigger an anthelminthic response.

## 2. Results

### Purified laminar layer and hydatid fluid from *Echinococcus granulosus* induce global translation

Dendritic cells were obtained from bone marrow of CF-1 mice and were culture for 6 days in complete medium supplemented with Flt-3-L as described in Methods. Prior to study translation levels in Bone Marrow Dendritic Cells (BMDCs), we analyzed by flow cytometry if *Eg* antigen stimulation *in vitro* induce cell viability loss. Cellular integrity was evaluated by propidium iodide incorporation. Figure 1A shows that neither pLL nor HF diminished BMDCs viability. The cell viability in the different assays was around 80-90%. BMDCs in presence of 3% of DMSO were used as a cell death control. We then analyzed whether pLL and/or HF were able to modulate global translation. To this purpose, we used the SUnSET method previously described by Schmidt *et al.* ^39^. Briefly, puromycin is a structural analog of aminoacyl tRNAs, which is incorporated into the nascent polypeptide chain and prevents elongation. When used in minimal amounts, puromycin incorporation in neosynthesized proteins reflects directly the rate of mRNA translation *in vitro* by immunodetection using an anti-puromycin antibody. In Figure 1B-C we tested the ability of 12D10 monoclonal antibody to monitor translation in single cells using immunofluorescence. As can be shown, both parasite compounds enhance translation in BMDCs compared to control cells. Specially, pLL induced a higher translation level compared to HF and showed statistically significant differences of puromycin detection, similar to that detected in LPS-stimulated BMDCs. As expected, cycloheximide, an inhibitor of eukaryotic translation elongation, or the anti-proliferative drug rapamycin displayed a dramatically reduced intensity of signal in treated cells. The expression of MHC class II protein was also evaluated in pLL- or HF-stimulated BMDCs and both *Eg* antigens did not show a statistically significant modulation. (Figure 1B-D).

**Figure 1.**
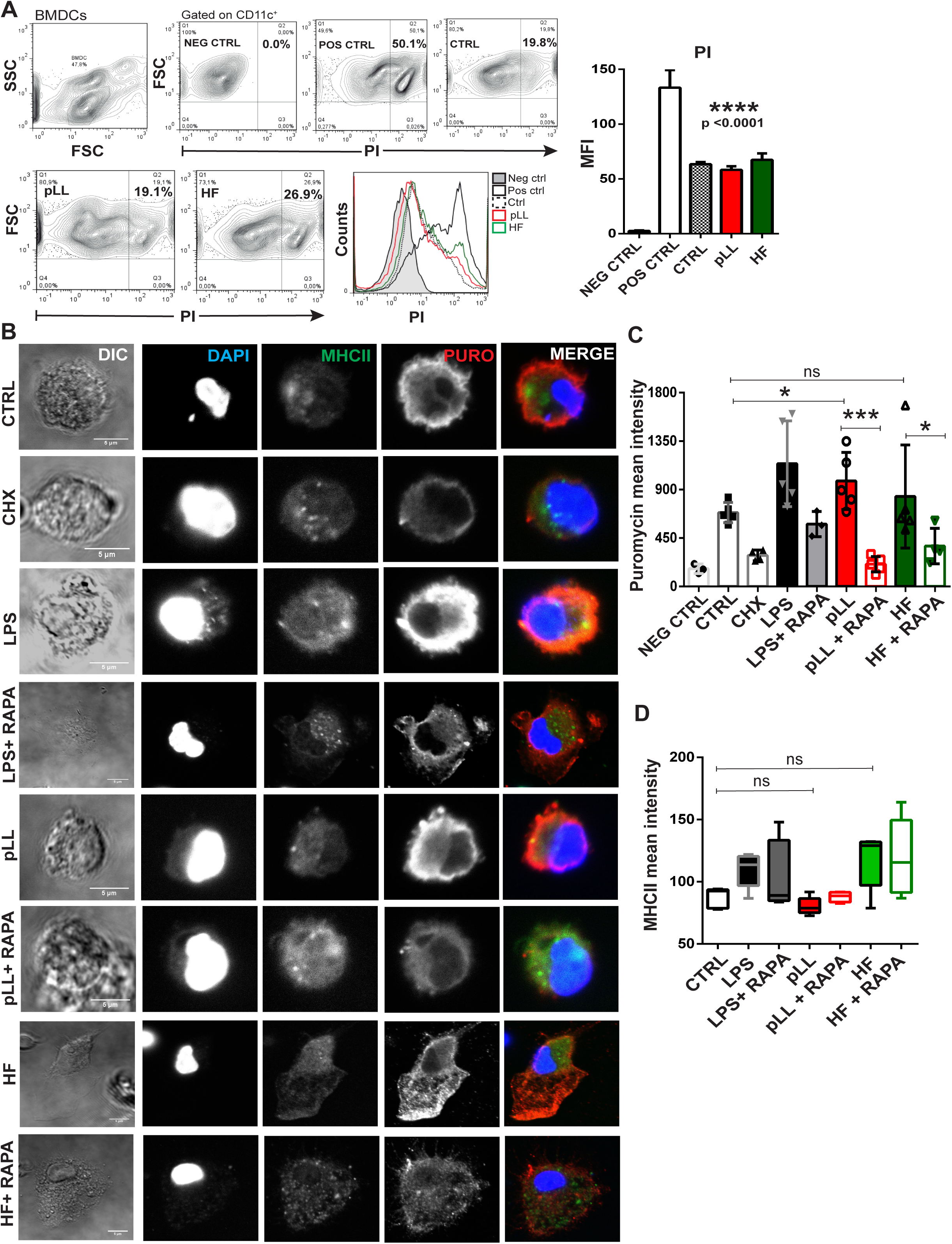
Global translation is induced by purified laminar layer and hydatid fluid from *Echinococus granulosus* in BMDCs. (A) BMDCs (1×10^6^/ml) were stimulated for 18h with pLL 20 μg or HF 5 μg from *E. granulosus.* Then, the viability of BMDCs was analyzed by flow cytometry using propidium iodide (PI). Positive control (POS CTRL) of necrosis represents BMDCs cultured in complete medium with 3% DMSO. Negative control (NEG CTRL) represents cells without PI staining and control (CTRL) shows BMDCs in absence of *Eg-*antigen stimulation. Dot blots of a representative experiment are shown (n=3). Data are expressed in percentage of positive cells or as the arithmetic means fluorescence intensity (MFI) ± SEM of n experiments (one-wayANOVA test ****p<0.0001) B) Protein synthesis was monitored in pLL- or HF-stimulated BMDCs by puromycin (1 μg/ml) incorporation after 18 h of *Eg* antigens stimulation. Immunofluorescence staining with the anti-puromycin mAb 12D10 was performed. Controls are cells not treated with puromycin (NEG CTRL), without any antigen-stimulation (CTRL), cells treated with cycloheximide (CHX) 5 min prior puromycin incorporation or LPS-treated cells (100 ng/ml). Rapamycin (20 nM) was added to stimulate-BMDCs when it is indicated. Scale bar, 5 µm. C) Bar graph show the puromycin mean intensity ± standard deviation of different cells in a representative experiment. (*p<0.05, ***p<0.001, pLL- or HF-stimulated cells vs controls; or vs rapamycin treatment when is indicated). Three independent experiments were performed. D) Bar graph show the MHC II mean intensity of different cells in a representative experiment (no significant differences were detected).

### Purified laminar layer and hydatid fluid of *Echinococcus granulosus* induce a phenotypic maturation and cytokines production unconventional by BMDCs

BMDCs phenotype was evaluated by flow cytometry. The purity of BMDCs was higher than 70% (range 65–80%) and their expression of CD11c, CD135, CD172a, CD370, CD205, MHC I, MHC II, CD86, CD80 and CD40 is shown in Figure 2A. In a set of experiments, we analyzed whether 20 μg of pLL or 5 μg of HF were able to modulate the expression of costimulatory molecules (CD86, CD40 and CD80) and the major histocompatibility molecules (MHC I and MHC II). After 18 h of culture, 1×10^6^/ml pLL- or HF-stimulated BMDCs were harvested from the culture and their phenotypes were analyzed by flow cytometry. LPS-stimulated BMDCs (100 ng/ml), cells cultured alone and in presence of 20 nM rapamycin were used as controls.

**Figure 2.**
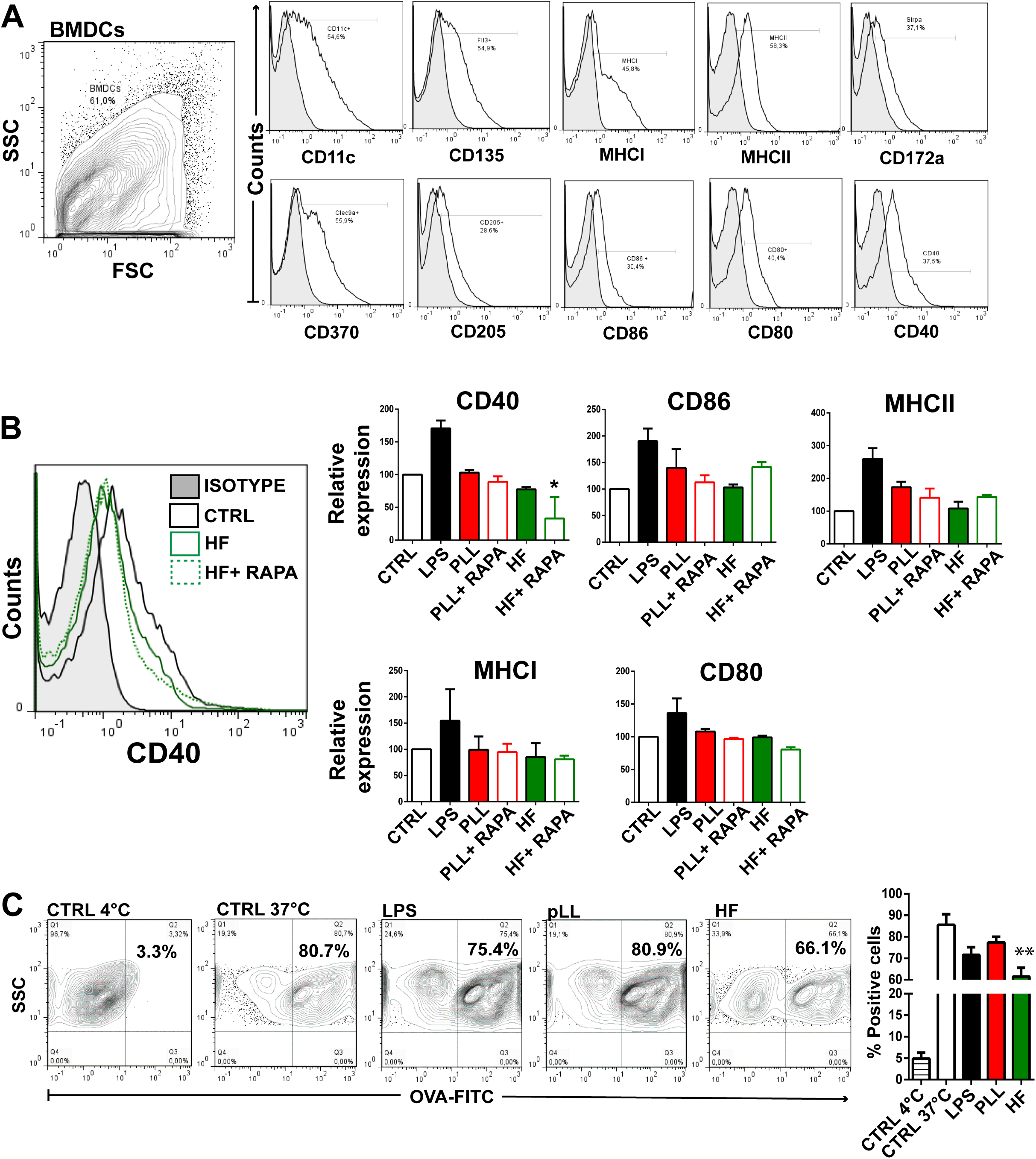
Purified laminar layer and hydatid fluid induce a slight maturation phenotype on BMDCs. BMDCs (1×10^6^/ml) were cultured alone (CTRL), with 20 μg pLL or 5 μg HF for 18 h. Then, BMDCs were harvested and their phenotype was analyzed by flow cytometry. (A) Histograms illustrating the phenotype of BMDCs at steady state based on the expression of CD11c, CD135, CD172a, CD370, CD205, MHC and co-stimulatory molecules. (B) Bar graphs show the relative mean fluorescence intensity (MFI) of MHC class I, MHC class II, CD86, CD80, and CD40 in the gate of CD11c^+^ cells for BMDCs cultured alone (white bars), with 100 ng/ml of LPS (black bars), in presence of pLL (red-filled bars), HF (green-filled bars) or rapamycin with *Eg* antigens (open bars). A representative histogram of CD40 expression is shown (n =4-6). Grey-filled histograms represent isotype controls (they were similar for BMDCs cultured alone or in the presence of *Eg* antigens). Results are the mean ± SEM of 4–6 experiments. (* p<0.05 vs BMDCs cultured alone). (C) Control BMDCs (4°C and 37°C) or pre-stimulated with 100 ng/ml of LPS, 5 μg of HF, or 20 μg of pLL were incubated with FITC-OVA (200 μg/ml) for 10 min at 37°C. The amount of endocytosis of FITC-OVA was measured by flow cytometry. A representative dot blot is shown. Results are expressed as a percentage of positive cells and represent the arithmetic mean of three experiments. The asterisks represent statistical significance (**p <0.01) for HF vs Control (37°C).

In our assays, BMDCs are shown as a unique cell population because the subpopulations analyzed: plasmacytoid dendritic cells (pDC, CD11c^+^ B220+ SiglecH^+^) and conventional dendritic cells (cDC1: CD11c^+^ CD24^+^, CD370^+^ CD172a^-^) and (cDC2: CD11c^+^, CD172a^+^ CD24^-^) did not show any differences in the response generated against the stimulation with both antigens used from *E. granulosus*. Figure 2B shows that pLL induced a non-statistically significant, but a trend in the up-regulation of CD86 and MHC II and that expression change was diminished by the use of rapamycin. Conversely, no changes in the expression of CD40, CD80 or MHC I were registered when BMDCs were stimulated with this acellular layer rich in polysaccharides. By contrast to pLL stimulation, a down-regulation of CD40 was observed in HF-stimulated BMDCs, and this modulation was statistically significant when rapamycin was present in the culture. Surprisingly, the effect of rapamycin in cultures with HF was uneven depending on the molecule studied. While MHC I and CD80 expression in membrane was slightly down-regulated, MHC II and specially CD86 expression showed a positive regulation. Additionally, to test if *Eg* macromolecules could modulate the uptake of antigen (Ag) by BMDCs we treated cells with pLL or HF and then incubated them with FITC-OVA, and analyzed their fluorescence by flow cytometry as a measure of Ag uptake ^40^. LPS was used as a control. This potent TLR4 agonist, showed in mature DCs a poor antigen uptake and migratory activity ^41^. The stimulation of BMDCs with *Eg* antigens during 18 h induced different rates of antigen-capture. On one hand, stimulation with pLL did not induce changes in endocytosis, but on the other hand, in the presence of HF, this activity markedly decreased compared to the control (Figure 2C).

Further studies were done to analyze whether pLL or HF were also able to stimulate the production of cytokines by BMDCs. To this aim, parasite antigens and BMDCs were cultured together for 18 h and the levels of TNF-α, IL-10, TGF-β, IL-12, IL-23, and IL-6 were measured by quantitative PCR. Figure 3 shows that BMDCs cultured alone express very low amounts of IL-6, TNF-α, IL-12, IL-23 and IL-10 but the expression of TGF-β was moderate, suggesting that they are in a resting state in a tolerogenic profile. As expected, treatment with LPS resulted in a marked stimulation in the production of cytokines (TNF-α, IL-6, IL-12, IL-10 and IL-23). When BMDCs were stimulated with pLL induce the expression of IL-6 and TNF-α, but not differences compared to control were observed in IL-12, IL-23, IL-10 and TGF-β. By contrast, only IL-6 and the anti-inflammatory cytokine IL-10 were up-regulated in BMDCs stimulated with HF. As expected, the use of the immunosuppressive drug rapamycin reduced gene expression in most of the cytokines studied.

**Figure 3:**
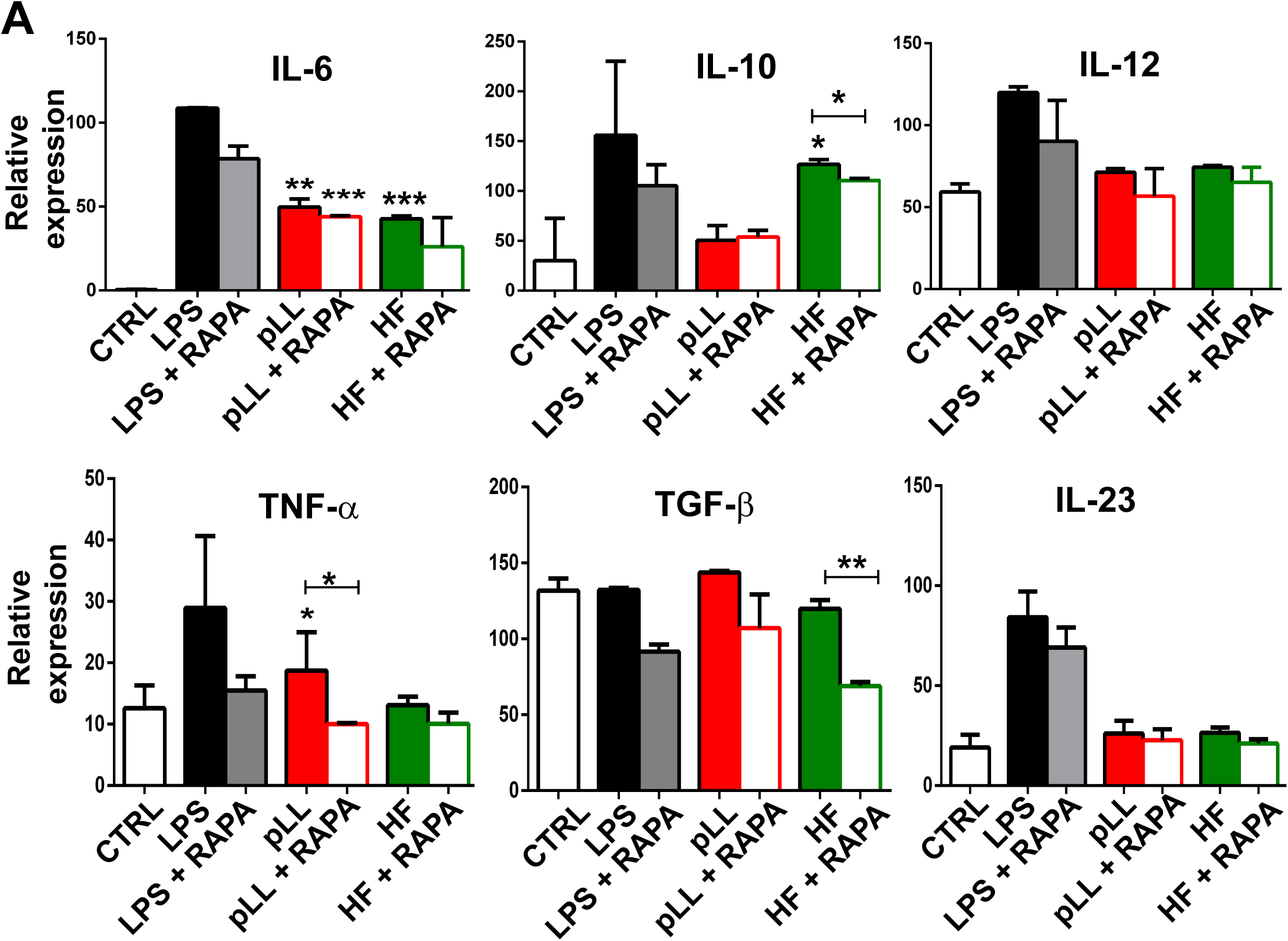
Purified laminar layer and hydatid fluid induce gene expression of pro- and anti-inflammatory cytokines by BMDCs. (A) BMDCs (1×10^6^/ml) were cultured alone (CTRL), with 100 ng/ml LPS, 20 μg pLL or 5 μg HF in presence or absence of 20 nM rapamycin for 18 h. Gene expression of different cytokines (IL-6, IL-10, IL-12, TNF-α, TGB-β and IL-23) was evaluated from isolated mRNA by quantitative PCR (relativized to the expression of GAPDH mRNA). Results are the mean ± SEM of three experiments performed in duplicate. (*p<0.05, **p<0.01, ***p<0.001 vs controls or when it is indicated stimulated-BMDCs vs rapamycin-treated BMDCs)

### *Echinococcus granulosus* antigens promote the capacity of dendritic cells to induce splenocyte proliferation *in vitro*

To evaluate the ability of BMDCs pre-stimulated with pLL or HF to function as antigen-presenting cells (APCs) and to promotes T cell proliferation, we performed a co-culture of BMDCs with CFSE labeled allogeneic splenocytes. Cell proliferation was analyzed by flow cytometry by measuring CFSE dilution. When splenocytes were cultured alone for 4 days, naïve T cells did not proliferate and more than 98% of the cells were CFSE^+^ (Figure 4). In contrast, when they were cultured together with LPS-stimulated BMDC for the same length of time, a large fraction of cells proliferated. Despite the phenotype described above and the profile of cytokines that pLL- and HF-stimulated BMDCs have shown, they have had high allostimulatory activity. Nevertheless, HF-stimulated BMDCs displayed more capacity to promote proliferation of splenocytes. In all conditions, when BMDCs were stimulated in the presence of rapamycin, the capacity of splenocytes proliferation induction was decreased in a range of 5% for pLL and 15% for HF.

**Figure 4:**
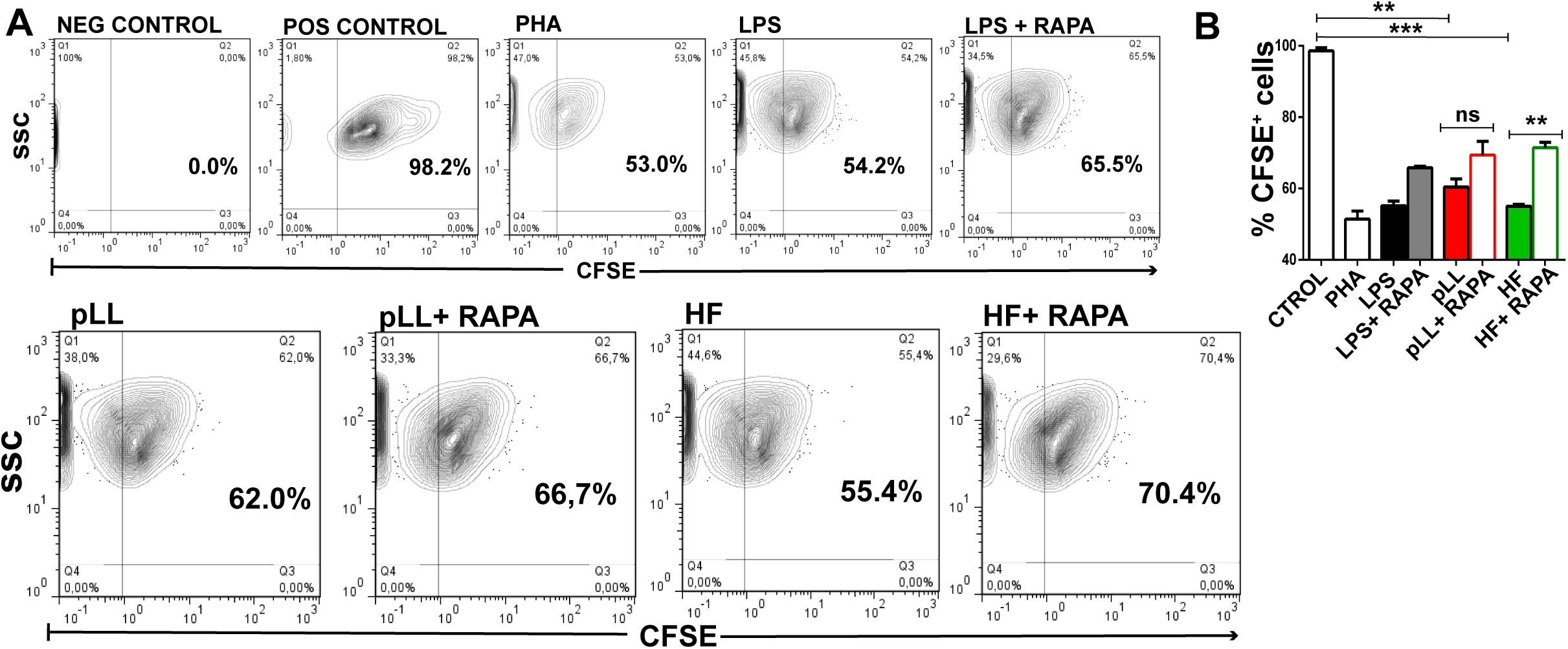
Purified laminar layer or hydatid fluid-stimulated BMDCs induce *in vitro* splenocyte proliferation. (A) Resting splenocytes were labeled with 1 μM CFSE and co-incubated with LPS, pLL or HF pre-stimulated BMDCs in presence or absence of rapamycin (20 nM) using a DC: splenocyte ratio of 1:4 and cultured in 96-well flat-bottomed plates in a final volume of 200 μl for 4 days. Splenocytes not labeled with CFSE (NEG CONTROL), without co-incubation with BMDCs (POS CONTROL) or treated with 1μg/ml of phytohaemagglutinin (PHA) were used as controls. Cells were harvested and analyzed for CFSE content by flow cytometry. A representative dot blot from one of three independent experiments, each done at least in duplicate is shown. (**p<0.01, ***p<0.001 vs controls or when it is indicated stimulated-BMDCs vs rapamycin-treated BMDCs).

### *Echinococcus granulosus* laminar layer activates the mTOR pathway in BMDCs

Different pathogens, including helminth parasites, modulate the mTOR pathway for controlling and shaping the effector responses of immune cells, through promoting or inhibiting host translation ^33,42^. Given that pLL- or HF-stimulated BMDCs enhances global translation, induces changes in their phenotype and cytokines expression, and effectively promotes T cell proliferation, we examined the activation of the mTOR pathway in this model. Thus, mTOR activated form (Ser2448) was immunodetected by confocal microscopy and western blot. It is known that this signaling pathway is modulated by growth factors, for this reason, the stimulation conditions were performed without the presence of FBS and Flt3L. As shown in Figure 5, mTOR activation levels were induced by both parasitic antigens. However, statistically significant differences were only observed in the presence of pLL. LPS stimulation or rapamycin treatment were used as controls. It has been previously shown that under favorable nutritional conditions the active mTORC1 is located on the endosomal membrane, instead, when there is nutritional stress, it is released to the cytoplasm ^43^. In our assays, we observed that the pattern of phosphorylated mTOR in HF-stimulated BMDCs was similar to that observed under control conditions, with a diffuse signal in the cytoplasm (Figure 5A). However, by immunoblot, the complex is in its active form as shown in Figure 5C. Otherwise, the presence of pLL induced a strong clustering of the signal near the cell nucleus, similar to that observed with LPS stimulation. Western blots also show a higher activation level of mTORC1 in pLL-stimulated BMDCs compared to HF-stimulated BMDCs.

**Figure 5.**
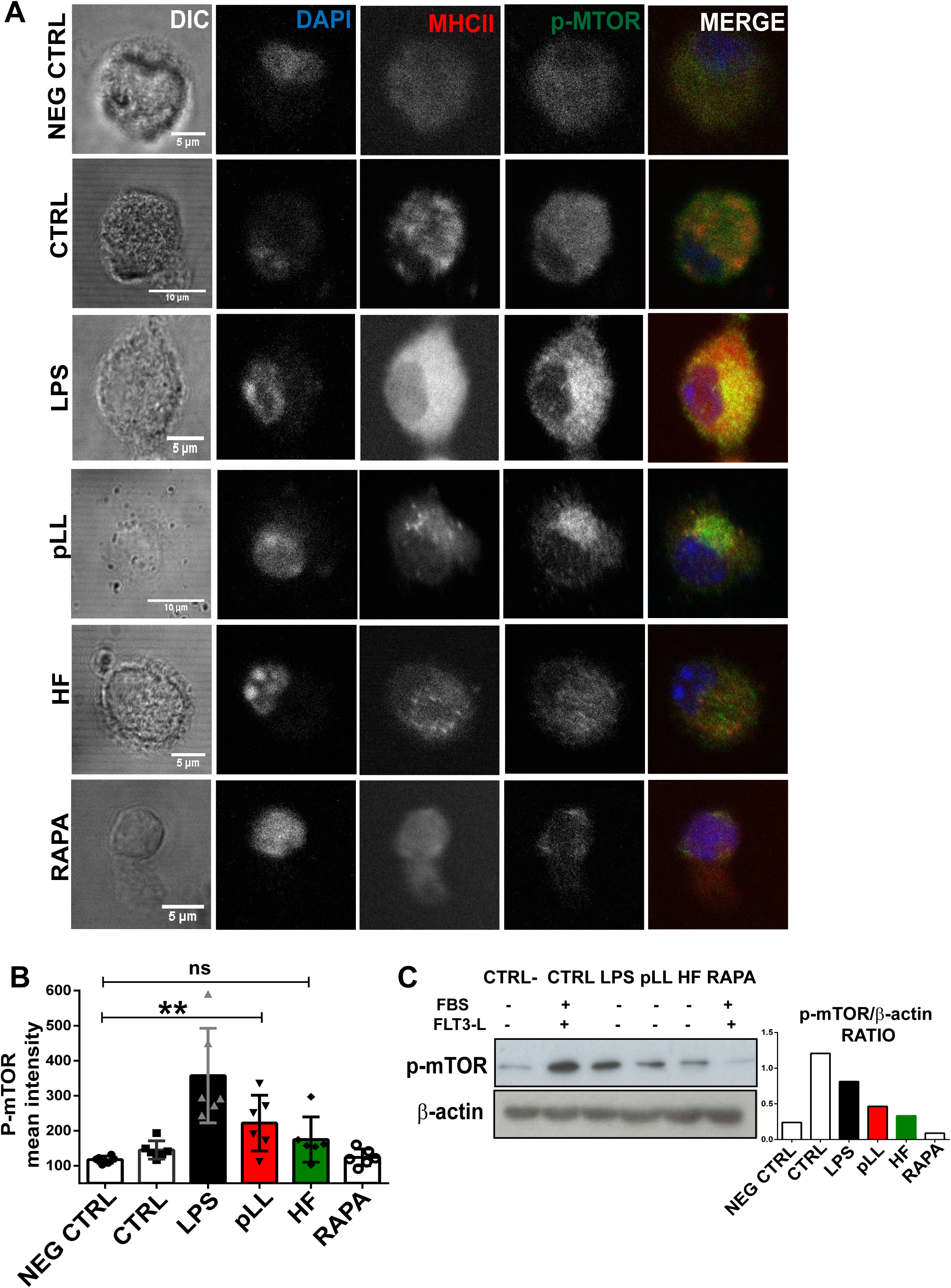
Purified laminar layer from *Echinococcus granulosus* induce the activation of mTOR complex. (A) BMDCs (1×10^6^/ml) were cultured in complete medium (CTRL), without FBS and Flt3L (NEG CTRL) or stimulated for 18h with 100 ng/ml LPS, 20 μg pLL or 5 μg HF in presence or absence of 20 nM rapamycin in growth factors depleted medium. Then, cells were stained with specific antibodies for MHC class II (red) and phosphor-mTOR (green) and visualized by confocal microscopy. Cell nuclei were stained with DAPI (blue). Scale bar 5 µm. (B) Bar graphs shows phosho-mTOR mean intensity ± standard deviation of different cells in a representative experiment of three independent experiments (**p<0.01 pLL-stimulated cells vs controls). (C) Immunoblot of phospho-mTOR is shown. β-actin was used for equal loading control. Phosphorylation signal was quantified with ImageJ software.

### Hydatid fluid stimulation induces autophagosome-like vesicles in BMDCs

Given that the autophagy is a basal process in DCs that contributes with the antigen presentation via MHC-II complexes in the resting state ^44^, we investigated its possible activation state upon antigen stimulation. Thus, in pLL- or HF-stimulated BMDCs, we immunostained LC3 on fixed cells and analyzed them by the immunofluorescence confocal microscopy. At normal conditions, LC3 is cleaved by a cysteine protease to produce LC3-I (18 kDa), which is located in the cytosolic fraction. However, upon exposure to various environmental stresses like starvation, LC3-I is converted to LC3-II (16 kDa). The latter is covalently attached to phosphatidylethanolamine on its C-terminus, and it binds tightly to autophagosome membranes. Therefore, LC3-II is considered the functional form of LC3 and has been used as a specific marker of autophagy ^45,46^. In Figure 6, BMDCs control cultures in presence of complete medium supplemented with Flt3L showed a cytoplasmic diffuse pattern of LC3, with a low number and small size of LC3-positive particles. As autophagy is known to be inhibited by serum-rich medium, autophagic structures in pLL- or HF-stimulated BMDCs were assessed in RPMI depleted of FBS and Flt3L at a density of 1×10^6^ cells/ml. HF-stimulated BMDCs significantly enhanced the mean fluorescence intensity of LC3 and showed a trend in the increment of number and the average size of LC3-positive structures in comparison with unstimulated cells cultured in the absence of growth factors (Figure 6A-B). Contrary, and in agreement with the higher-level of phosphorylated mTOR observed before, pLL-stimulated BMDCs exhibited fewer LC3-positive particles compared to the control condition. Cell culture in presence of Hank’s Balanced Salt Solution and rapamycin-treated cells were used as a positive control of occurrence of autophagocytic process.

**Figure 6.**
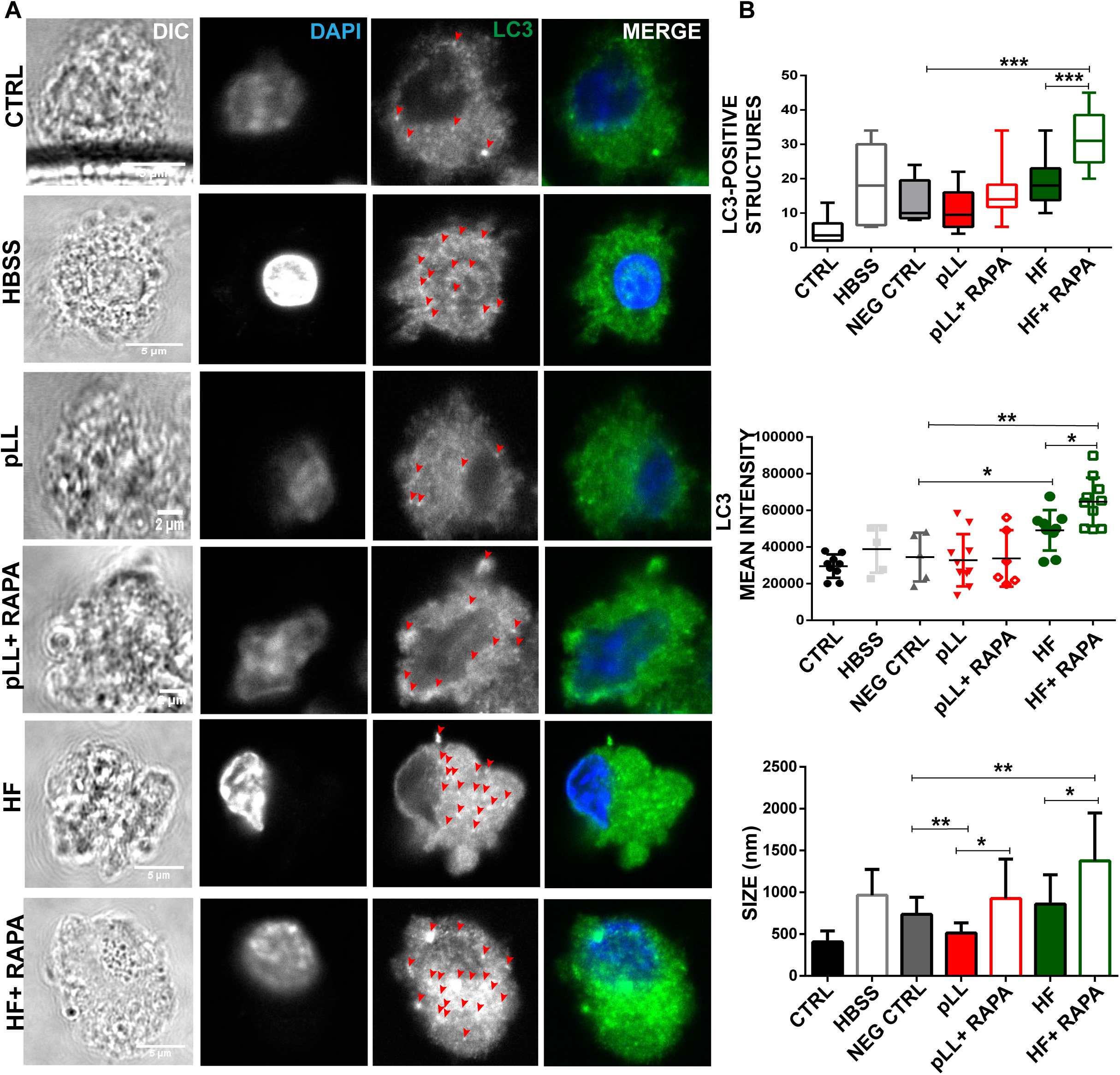
Hydatid fluid enhances the number of LC3-positive structures in BMDCs. Confocal images of BMDCs (1×10^6/^ml) stimulated during 18h with pLL or HF from *E. granulosus* in presence or absence of (20 nM) rapamycin. Cells culture without stimulation (CTRL), in starvation condition (HBSS) and without grown factors (NEG CTRL) were used as controls. Cells were stained with specific antibody for LC3 (green), and cell nuclei were stained with DAPI (blue). Scale bar 5 µm. Arrows indicate positive LC3 dotted structures in the cell cytoplasm. (B) Bar graphs show number, MFI and size of LC3^+^ structures ± standard deviation of different cells in a representative experiment of three independent experiments (*p<0.05, **p<0.01, ***p<0.001 pLL or HF-stimulated cells vs controls).

## 3. Discussion

In this report, we provide new evidence about the link of immunological pathways of parasitic antigens detection in DCs, with the metabolic signaling pathways that govern global translation, growth, proliferation, survival, and autophagy. We demonstrated how DCs through mTORC1 adapt their transcriptional responses and protein translation to control phenotype maturation, cytokine production and T cell proliferation under *Eg* antigen stimulation.

Previous publications have reported the induction of apoptosis by cestode excretory/secretory-products in DCs ^47^, macrophages ^48^, and HF-treated lymphocytes ^49^. In our assays, we corroborated that the stimulation of BMDCs with pLL and HF did not cause a significant decrease in cell viability (Figure 1A), thus both antigens were used for the subsequent studies. Recognition of parasite-derived molecules induced not only the production of specific genes like cytokines, co-stimulatory molecules, or the up-regulation of MHC; but also promoted the transcription and translation of hundreds of genes, in order to control infection ^50,51^. In this sense, mTOR appears to be part of the early sensing machinery, bridging the gap between PRR signaling and the cellular metabolism required to organize and trigger an effective immune response. There are no previous data about the levels of host protein neosynthesis and the signaling pathways implicated after stimulation with *Eg* antigens. As has previously been described in other pathogens ^52,53^, pLL and HF *Eg* antigens significantly promoted overall protein synthesis compared to untreated cells, and this effect was shut off by the use of rapamycin (Figure 1B), an inhibitor of mTORC1. In line with these results and in the absence of growth factors, both parasitic antigens stimulated phosphorylation of mTOR (Ser2448) in BMDCs, being this effect higher with pLL compared to HF (Figure 5). Previous studies in *E. granulosus* have shown in GM-CSF derived DCs or M-CSF derived macrophages that exposure to pLL strongly inhibited PI3K, Akt and GSK3 phosphorylation induced by LPS, and that stimulation of DCs with pLL alone inhibited basal phosphorylation of these proteins ^37,38^. On the other hand, *Eg* excretory/secretory products induced alternative activated macrophages phenotype (M2), through the activation of the PI3K/AKT/mTOR pathway ^36^. Our data are in line with those reported by Wang *et al.* ^36^, but are not contrary to those described by Pittini *et al.* ^37^, given that the presence of pLL did not inhibit Akt or GSK3 phosphorylation in Flt3L-BMDCs.

Modulation of co-stimulatory and MHC molecules in DCs are linked to mTOR activation ^12,19,22,54^. A decrease in the upregulation of MHC and co-stimulatory molecules in BMDCs have been previously described in presence of rapamycin ^55^. We have observed that pLL-stimulated BMDCs displayed a partial maturation phenotype with a slight increase in CD86 and MHC II reverted by rapamycin. In contrast, in presence of HF, only a decrease in CD40 was observed, which was also enhanced by this immunosuppressive drug (Figure 2B). This slight phenotypic maturation in DCs could be explained by the absence of activation in the NFκB and MAP kinases pathways ^38^. We have previously reported that exosome-like vesicles induce maturation of BMDCs with an increase of CD86 and a slight down-regulation in the expression of MHC II molecule ^56^. Likewise, previous reports have shown a weak maturation phenotype in mouse and human DCs upon stimulation with *Eg* antigens in combination with potent immunogens ^36,38,57,57–61^. Uptake and processing of antigens by immature DCs are critical steps in producing an immune response. Thus, any treatment that induces maturation in BMDCs will decrease their antigen-capture capacity. In our assays, treatment with HF but not with pLL induced a diminished uptake of antigens (Figure 2C). Similar results were observed during DC differentiation, where in presence of *Eg* antigens the capacity to take up dextran has been markedly reduced ^62^. The mTORC1 activation in pLL-stimulated BMDCs not only raise antigen presentation by phenotype maturation, but also begin to induce an early response with inflammatory cytokines such as IL-6 and TNF-α (Figure 3). Subsequently, the release of protein antigens that are presents in HF induces the expression towards immunomodulatory cytokines such as IL-10 in a mTORC1-dependent manner (Figure 3). These results strongly confirm the events supported by Sukhbaatar *et al*, referred to the function of mTOR to modulate activation profile in DCs and support T cell stimulation or tolerance in the surrounding tissue. ^54^. In this context, different parasites have evolved to regulate mTOR signaling to influence DCs and macrophages function. In the helminth *B. malayi*, it has been reported the mTORC1 inhibition in DCs through secreted rapamycin-like molecules, which induced a significant downregulation of p70S6K and 4EBP1 phosphorylation and impaired the synthesis of IL-12 and IL-10 reducing their CD4^+^ T cell-activating capacity ^33^. On the other hand, and in concordance with this work, *Toxoplasma gondii*-infected macrophages induced the mTORC1 activation increasing translation of pro-inflammatory cytokines such as IL-6, and also cytokines as IL-10, with anti-inflammatory feature ^63^.

Like other helminth infections, echinococcosis induces an acute Th1/Th17 and a chronic Th2 cytokine profile patterns. Remarkably, in our experiments, pLL-stimulated BMDCs induced the expression of pro-inflammatory cytokines (Figure 3). In the same line of evidence, it has been described that pLL-stimulated DCs induces the production of the pro-inflammatory IL 1-β through the activation of the NLRP3 inflammasome, suggesting that the presence of inflammation contributes to Th1 and Th17 responses ^64^. IL-6 could be involved in favoring Th17 profile and promoting host susceptibility by limiting protective Th2 responsiveness. However, a local inflammatory environment, in the presence of IL-6 and IL-1β, can promote tolerance by inducing IL-10-producing regulatory B cells ^65^. In our case, the anti-inflammatory cytokine IL-10, was only detected in the presence of HF. Taken together, several reports have demonstrated that mTOR specially supports Th2 lymphocyte activation since it promotes IL-10 production by DCs ^66^. Particularly, this regulatory cytokine is abundantly expressed by leukocytes in infected hosts, especially in the immediate vicinity of the parasite ^5^. IL-10 secreted by HF-stimulated DCs could promote T cell differentiation to Treg1, specialized in immunosuppressive capacity through the production of high amounts of IL-10 ^67^. Immunopathology of alveolar and cystic echinococcosis revealed the presence of CD4^+^ CD25^+^ Treg cells producing IL-10 and TGF-β ^68,69^. Gene expression of IL-12, TGF-β, and IL-23 in *Eg*-antigen stimulated DCs did not show changes compared to untreated cells (Figure 3). In all conditions tested, rapamycin diminished gene transcription. HF and particularly Antigen B reduced the production of IL-12p70 and TNF-α in LPS-stimulated DCs ^60,61^. The same expression pattern was observed when DCs were exposed with HF during differentiation. HF impaired its subsequent ability to secrete IL-12, IL-6, or PGE2 in response to stimulation with LPS. Nevertheless, HF can stimulate maturation by the release of IL-12 and IL-6 in differentiated DCs ^59^. Excretory/secretory products of *Echinococcus* failed to induce the secretion of pro-inflammatory IL-12p70 in DCs, but the production of immunosuppressive IL-10 was enhanced ^47^. In response to HF, DCs produced IL-6, but not IL-12 or IL-10, nevertheless, these cytokines were produced at high levels when DCs were stimulated with Eg-ferritin ^70^. Similar to DC maturation profile, *Eg* antigens stimulation induced a slight expression of cytokines, which must be analyzed in co-stimulation with a strong TLR agonist.

Although differences in the antigen capture, phenotypic maturation and cytokine expression were observed between HF- and pLL-stimulated BMDCs, both parasitic antigens were strong promoters of *in vitro* splenocyte proliferation and this capacity was rapamycin-sensitive (Figure 4). Previous data showed a specific inhibitory effect of pLL on macrophages proliferation in response to IL-4 or M-CSF, but increased antigen-specific activation of T cells have been observed ^37^. Likewise, a high capacity of *Echinococcus-*antigen pulsed DCs to induce allostimulatory T cell proliferation has been previously described ^62^. Moreover, mice infected with *Echinococcus* for 4 months have shown increased T cell proliferative response to ConA ^71^

Interestingly, it is known that mTOR attenuates autophagy-dependent presentation of endogenous antigens, but after pathogen recognition, it promotes the presentation of exogenous antigens by favoring the lysosome acidification and the MHC-II/peptide traffic to the cell membrane ^54,66^. Differences in mTORC1 activation and phenotypic maturation led us to wonder what levels of autophagic processes occurred in DCs with both antigenic stimulations. We report for the first time that the *Echinococcus* HF-stimulated DCs induced an increase in the autophagic process (Figure 6). This observation, correlates with the lower level of mTOR activation in this condition (Figure 5). Similarly, associated to mTORC1 inhibition, *B. malayi*-stimulated DCs have been described an increase of autophagy in these cells by upregulation of phosphorylated Beclin 1 and degradation of p62 ^33^.

Therefore, we can conclude that to orchestrate a specific anti-parasite response, DCs regulates cytokine and co-stimulatory molecules expression, promoting T-cell proliferation through the link of two main signaling pathways, the innate recognition of antigens and the master regulator of nutrient-sensing. A better understanding of these early events is needed to develop rational strategies that prevent the establishment of the parasite in host tissue.

## 4. Methods

### Ethics statement

The experimental protocol was approved by the Animal Care and Use Committee at the Faculty of Exact and Natural Sciences, Mar del Plata University, Argentina. The mice were handled in agreement with National Health Service and Food Quality (SENASA) guidelines, Argentina and with The Guide for the Care and Use of Laboratory Animals published by NIH, U.S. (2011) https://grants.nih.gov/grants/olaw/guide-for-the-care-and-use-of-laboratory-animals.pdf.

### Mice

Female CF-1 mice 5-7 weeks old and weighed 28–35 g were provided by the SENASA. *E. granulosus* cysts were obtained from the peritoneal cavity of mice after 6-8 months of injected with 1500 protoscoleces in suspension. The behavior of the mice was observed daily for any signs of discomfort or distress (ruffled coat, hunched posture, lethargy). All efforts were made to minimize suffering. Infected mice were anesthetized with ketamine-xylazine (50 mg/kg/mouse-5 mg/kg/mouse) and sacrificed by cervical dislocation. Then, the samples were collected post-mortem.

### Purification of hydatid fluid and laminar layer of *Echinococcus granulosus*

Hydatid cysts were collected aseptically from infected cattle slaughtered. HF was punctured from the cysts and then centrifuged at 2000 g for 10 min at 4°C. The filtred supernatant was conserved and protein concentration was determined by absorbance at 280 nm. HF was free of Mycoplasma as measured by the Mycoplasma PCR detection kit (Venor®GeM). LL was isolated from cysts obtained from infected mice and was purified by following the described instruction in Seoane *et al*. ^37^.

### Generation of Bone Marrow derived Dendritic Cells

BMDCs were obtained by flushing bone marrow of femurs and tibias of CF-1 mice as previously described with minor modifications ^72^. Cells were plated at 1×10^6^/ml with RPMI 1640 supplemented with 5% heat-inactivated fetal bovine serum, 100 U/ml penicillin/streptomycin, 10 μg/ml gentamicin and 2 mM L-glutamine, (all from Thermo Fisher) and in presence of 100 ng/ml Flt3L (R&DSystems) at 37°C in 5% CO_2_ for 6 days. Finally, DC-population was characterized by flow cytometry using fluorescence-conjugated monoclonal antibodies (mAb) directed against CD11c (HL3), Flt3 (A2F10), Clec9a (42D2), CD172a (P84), CD11b (M1/70), CD3 (145-2C11), CD45R/B220 (RA3-6B2), SiglecH (eBio440c), and CD24 (M1/69) (eBiosciences). Approximately 70–80% of the cells were CD11c^+^.

### Confocal microscopy

For immunofluorescence staining, BMDCs were harvested after stimulation with 100 ng/ml LPS, 20 μg pLL or 5 μg of HF in presence or absence of 20 mM rapamycin, and seeded on alcian blue-treated coverslips during 20 min. Then, the cells were washed and blocked with PBS-BSA 2% in a wet chamber for 30 min, fixed in 4% PFA for 10 min on ice, washed twice with 0.1 mM glycine in PBS and permeabilized with 0.05% saponin. Afterward, they were incubated with mAb MHC class II-FITC (1:100, clone M5 / 114.15.2), anti-puromycin (1:100, clone 12D10), Phospho-mTOR (Ser2448), (1:100, clone D9C2) or LC3-β, (1:100, clone H50) overnight at 4°C. After washings in PBS 0.1% Tween-20, cells were incubated with rabbit anti-mouse secondary antibody conjugated with Alexa 555 (1:400, A-21427) or Alexa 488 (1:400, A-11059) and with donkey anti-rabbit Alexa 594 (1:400, R37119) for 1 h at 37 °C. Then the cells were washed and incubated with 50 ng/ml DAPI (Sigma-Aldrich) to counterstain nuclei. Coverslips were mounted on glass slides using Fluoromount G. Immunofluorescence images were acquired with an inverted confocal laser scanning microscope (Nikon, Confocal Microscope C1) using a 60 x oil immersion objective. Image analysis was performed using Fiji software, in which background was reduced using brightness and contrast adjustments applied to the whole image. Fluorescent intensity graphs were performed using the Histogram plugin in Fiji software. Briefly, to quantify MHC II, global translation, or phosphorylation levels of mTORC1, a total of ten cells in all conditions were analyzed. Image files were loaded as separate image stacks. LC3-positive intracellular compartments were manually quantified.

### Quantitative reverse transcription polymerase chain reaction

Total RNA was isolated from stimulated-DCs cultured 18h using TRIzol® Invitrogen, according to the manufacturer’s instructions. The quantity and purity of the isolated RNA were evaluated using a NanoDrop ND-1000 Spectrophotometer. 100 ng to 1 µg total RNA was subjected to reverse transcription using random primers and M-MLV RT (Invitrogen). Specific primer pairs for each gene were designed: IL-10 (fw: CCAAGCCTTATCGGAAATGA; rev: TTTTCACAGGGGAGAAATCG), TGF-β (fw: TTGCTTCAGCTCCACAGAGA; rev: TGGTTGTAGAGGGCAAGGAC), IL-6 (fw: AGTTGCCTTCTTGGGACTGA; rev: TCCACGATTTCCCAGAGAAC), TNF-α (fw: AGCCCCCAGTCTGTATCCTT; rev: CTCCCTTTGCAGAACTCAGG), IDO (fw: GGCTAGAAATCTGCCTGTGC; rev: AGAGCTCGCAGTAGGGAACA), IL-12p35 (fw: CATCGATGAGCTGATGCAGT; rev: CAGATAGCCCATCACCCTGT), IL-23p19 (fw: GACTCAGCCAACTCCTCCAG; rev: GGCACTAAGGGCTCAGTCAG). Gene expression analysis was performed on a 7500 Real Time PCR System (Applied Biosystems) using SYBR®Green PCR Master Mix for detection of PCR product (Applied Biosystems). PCR assays were carried out under the following conditions: a holding stage of 95°C (10min), a cycling of 40 steps of 95°C (15s), 60°C (1 min), plus a melting curve stage of 95°C (15s), 60°C (1 min), 95°C (15s). Expression level was evaluated using the 2^-ΔΔCt^ method, and each experiment was performed in duplicate with appropriate non-template controls. The relative amount of each transcript was determined by normalizing to GAPDH.

### Flow cytometry

BMDCs were harvested, washed with PBS with 2% FBS and suspended in a solution containing fluorescein isothiocyanate (FITC), phycoerythrin (PE) and phycoerythrin cyanine 5 (PECy5) - conjugated mAbs at saturating concentrations for 30 min at 4°C. Two additional washes were then performed to eliminate no bound antibodies. mAb directed to CD11c (HL3), CD24 (M1/69), CD45R/B220 (RA3-6B2), SiglecH (eBio440c), CD40 (HM40-3), CD80 (16-10A1), CD86 (GL1), MHC class I (AF6-88.5.5.3) and MHC class II (M5 / 114.15.2) CD135 (A2F10), CD370 (42D2), CD205 (205yekta) were from eBioscience and CD172a (P84) was from BioLegend. In all cases, isotype-matched control antibodies were used, and a gate (R1) was defined in the analysis to exclude all nonviable cells and debris, based on size and PI staining. The analysis was performed using a Partec Cyflow Space flow cytometer, and the FlowJo software. The results are expressed as the mean fluorescence intensity or as the percentage of positive cells.

### Immunoblotting

Total proteins isolated from BMDCs were lysed in CytoBuster protein extraction reagent (Novagen). Protein quantification was performed using NanoDrop One Spectrophotometer. 30 μg of proteins were loaded for all samples and analyzed by SDS-PAGE, transferred onto a nitrocellulose membrane (Amersham) at 120 mA for 60 min. Following, the membranes were incubated in blocking solution (TBST: 20 mM Tris-HCl, 150 mM NaCl, 1% Tween-20, pH 7.6 containing 2% bovine serum albumin for 4 h at 20°C) and were probed with a 1:1000 dilution of rabbit polyclonal antibody raised against mouse Phospho-mTOR (Ser2448) and stained with a secondary peroxidase-conjugated goat anti-rabbit IgG. As load control, the presence of β-actin was revealed (clone 8H10D10) Development was performed with ECL Western blotting reagents (GE Healthcare, UK).

### Translation intensity measurement

Puromycin labeling for measuring the intensity of translation was performed as previously described.^73^ For confocal microscopy, 1 μg/mL puromycin (Sigma), diluted in PBS was added to the culture medium 15 min before harvesting at 37°C and 5% CO_2_. Where indicated, 10 μM cycloheximide (Sigma) was added 5 min before puromycin. Cells were then harvested and seeded on coverslips, as previously explained. Next, puromycin incorporation was revealed by the 12D10 antibody. Global neo-synthetized proteins were evaluated in stimulated-BMDCs with or without rapamycin 20nM for 18 h. Cells not treated with puromycin and 100ng/ml LPS-activated BMDCs were used as controls.

### Antigen uptake by BMDCs

Antigen uptake was measured in BMDCs using a standard method ^40^. FITC labeled-chicken ovalbumin (FITC-OVA) was used as the antigen. Briefly, BMDCs were cultured in untreated conditions or treated with 100ng/ml LPS, 5μg of HF, or 20μg of pLL for 18 h. Cells were then washed with medium prior to the addition of FITC-OVA at a final concentration of 200 μg/ml in a final volume of 200 μl of RPMI 1640 medium containing 10% FCS for 10 min at 37°C. After washed with cold PBS, containing 0.5% BSA, the cells were suspended in PBS for the flow cytometry analysis. Cells without antigen or incubated with OVA-FITC at 4°C to inhibit intracellular uptake were used as negative controls.

### *In vitro* proliferation assay

Stimulated-BMDCs were washed, counted, and replated for incubation with freshly isolated splenocytes. The spleens were harvested from euthanized CF-1 mice and placed into 10 ml of RPMI 1640 supplemented with 5% heat-inactivated fetal bovine serum, 100 U/ml penicillin/streptomycin, 10 μg/ml gentamicin and 2 mM L-glutamine, (all from Thermo Fisher). The spleens were prepared into a single-cell suspension using a laboratory homogenizer, washed twice, and stained with the 5-carboxyfluorescein diacetate succinimide ester (CFSE) at a concentration of 1 μM, according to the manufacturer’s instructions. CFSE-stained splenocytes were co-incubated with BMDCs using a DC: splenocyte ratio of 1:4 (25×10^4^ BMDCs / 1×10^5^ splenocytes) and cultured in 96-well flat-bottomed plates in 200 μl of RPMI 1640 supplemented with 0.1% FCS. Cellular proliferation was assayed after 4 days of culture by flow cytometry in total splenocytes population. Splenocytes without labeling with CFSE, without co-incubation with BMDCs or treated with 1 μg/ml of phytohaemagglutinin (PHA) were used as controls.

### Statistical analysis

Statistical analysis was performed using GraphPad Prism Software. The most appropriate statistical test was chosen according to the set of data. One-way ANOVA, Student’s T-test and Mann Whitney test were used. *p<0.05, **p<0.01, ***p<0.001, ****p<0.0001.

## 6. Acknowledgements

The authors acknowledge all FCEyN cytometry and Imaging core facilities for expert assistance. Specially, Lic. Daniela Villamonte (Servicio de Microscopía Confocal, CONICET, Universidad Nacional de Mar del Plata), Lic. Viviana Daniel (Servicio de Citometría de Flujo, CONICET, Universidad Nacional de Mar del Plata). We also thank Dra. Alejandra Goya (SENASA, Argentina). The laboratory is supported by grants from National University of Mar del Plata, Ministry of Science, Technology and Productive Innovation, and CONICET. The authors declare to have no competing interests.

## 7. Authors’ Contribution

C.R.R., M.C.N., M.C., M.M., N.P, J.L. performed research. M.C., A.C.C., C.R.R designed research and analyzed data. A.C.C., C.R.R. wrote the paper.

## 8. Competing interests

The authors declare no competing interests.

## Notes

### Competing Interest Statement

The authors have declared no competing interest.

